# Age-Dependent Formation of TMEM106B Amyloid Filaments in Human Brain

**DOI:** 10.1101/2021.11.09.467923

**Authors:** Manuel Schweighauser, Diana Arseni, Melissa Huang, Sofia Lövestam, Yang Shi, Yang Yang, Wenjuan Zhang, Abhay Kotecha, Holly J. Garringer, Ruben Vidal, Grace I. Hallinan, Kathy L. Newell, Airi Tarutani, Shigeo Murayama, Masayuki Miyazaki, Yuko Saito, Mari Yoshida, Kazuko Hasegawa, Tammaryn Lashley, Tamas Revesz, Gabor G. Kovacs, John van Swieten, Masaki Takao, Masato Hasegawa, Bernardino Ghetti, Benjamin Falcon, Alexey G. Murzin, Michel Goedert, Sjors H.W. Scheres

**Affiliations:** Medical Research Council Laboratory of Molecular Biology, Cambridge, UK; Thermo Fisher Scientific, Eindhoven, The Netherlands; Department of Pathology and Laboratory Medicine, Indiana University School of Medicine, Indianapolis, IN, USA; Department of Brain and Neurosciences, Tokyo Metropolitan Institute of Medical Science, Tokyo, Japan; Molecular Research Center for Children’s Mental Development, United Graduate School of Child Development, University of Osaka, Osaka, Japan; Department of Neurology, National Center Hospital, National Center of Neurology and Psychiatry, Tokyo, Japan; Department of Neuropathology, Tokyo Metropolitan Geriatric Hospital and Institute of Gerontology, Tokyo, Japan; Institute for Medical Science of Aging, Aichi Medical University, Nagakute, Japan; Division of Neurology, Sagamihara National Hospital, Sagamihara, Japan; Department of Neurodegenerative Disease and Queen Square Brain Bank for Neurological Disorders, UCL Queen Square Institute of Neurology, London, UK; Tanz Centre for Research in Neurodegenerative Diseases and Department of Laboratory Medicine and Pathobiology, University of Toronto, Toronto, Canada; Institute of Neurology, Medical University of Vienna, Vienna, Austria; Department of Neurology, Erasmus Medical Centre, Rotterdam, The Netherlands; Department of Clinical Laboratory, National Center of Neurology and Psychiatry, National Center Hospital, Tokyo, Japan; Department of Neurology, Mihara Memorial Hospital, Isesaki, Japan

**Author notes:** These authors contributed equally; except for M.S. they are in alphabetical order. These authors jointly supervised. Correspondence to and. Medical Research Council Prion Unit, Institute of Prion Diseases, University College London, UK.

## Abstract

Many age-dependent neurodegenerative diseases, like Alzheimer’s and Parkinson’s, are characterised by abundant inclusions of amyloid filaments. Filamentous inclusions of the proteins tau, amyloid-β (Aβ), α-synuclein and TDP-43 are the most common. Here, we used electron cryo-microscopy (cryo-EM) structure determination to show that residues 120-254 of the lysosomal type II transmembrane protein 106B (TMEM106B) also form amyloid filaments in the human brain. We solved cryo-EM structures of TMEM106B filaments from the brains of 22 individuals with neurodegenerative conditions, including sporadic and inherited tauopathies, Aβ-amyloidoses, synucleinopathies and TDP-43opathies, as well as from the brains of two neurologically normal individuals. We observed three different TMEM106B folds, with no clear relationship between folds and diseases. The presence of TMEM106B filaments correlated with that of a 29 kDa sarkosyl-insoluble fragment of the protein on Western blots. The presence of TMEM106B filaments in the brains of older, but not younger, neurologically normal individuals indicates that they form in an age-dependent manner.

## Introduction

TMEM106B is a type II transmembrane protein of 274 residues that localises to late endosomes and lysosomes (Nicholson and Rademakers, 2016; Fen et al., 2021). It is expressed ubiquitously, with highest levels in brain, heart, thyroid, adrenal and testis (www.proteinatlas.org; Uhlén et al, 2015). Reminiscent of the amyloid precursor protein APP, TMEM106B is sequentially processed through ectodomain shedding, followed by intramembrane proteolysis, with possible variability in the intramembrane cleavage site. Lysosomal proteases have been implicated in the cleavage of TMEM106B in the C-terminal lumenal domain, but no specific enzymes have been identified. Although the cleavage site is unknown, it has been shown indirectly to be at a position close to glycine residue 127. The resulting C-terminal fragment contains five glycosylation sites at asparagine residues 145, 151, 164, 183 and 256. Following shedding of the ectodomain, the N-terminal fragment is cleaved by signal peptide peptidase-like 2a (SPPL2a), possibly at two different sites around residue 106 (Brady et al., 2014).

Genetic variation at the *TMEM106B* locus has been identified as a risk factor for frontotemporal lobar degeneration with TDP-43 inclusions (FTLD-TDP), especially for individuals with granulin gene (*GRN*) mutations (Van Deerlin et al., 2010). Levels of TMEM106B are elevated in FTLD-TDP (Chen-Plotkin et al., 2012). TMEM106B has also been reported to be involved in other diseases (Nicholson and Rademakers, 2016; Fen et al., 2021). Genome-wide association studies have implicated *TMEM106B* with variability in age-associated phenotypes in the cerebral cortex (Rhinn and Abeliovich, 2017).

Previously, cryo-EM imaging allowed atomic structure determination of filaments of the proteins tau (Fitzpatrick et al., 2017; Falcon et al., 2018; Falcon et al., 2019; Zhang et al., 2020; Shi et al., 2021), α-synuclein (Schweighauser et al., 2020), and Aβ (Kollmer et al., 2019; Yang et al., 2021) that were extracted from the brains of individuals with different neurodegenerative diseases. The cryo-EM structures revealed that distinct folds characterise different diseases. For tauopathies, this has made it possible to classify known diseases further and to identify new disease entities (Shi et al., 2021).

Cryo-EM structure determination can also be used to identify previously unknown filaments. Here, we have used cryo-EM to show that residues 120-254 from the lumenal domain of TMEM106B form amyloid filaments in the human brain. We initially observed TMEM106B filaments in the brains of individuals with familial and sporadic tauopathies, Aβ-amyloidoses, synucleinopathies and TDP-43opathies. However, the role of TMEM106B filaments in disease remains unclear. They were not observed in brains from young individuals, but their presence in brains from older, neurologically normal controls indicates that TMEM106B filaments may form in an age-dependent manner.

## Results

Using sarkosyl extraction protocols that were originally developed for α-synuclein (Tarutani et al., 2018, Schweighauser et al, 2020), we observed a common type of filament that lacked a fuzzy coat in the cryo-EM micrographs from cases of various neurodegenerative conditions. Structure determination to resolutions sufficient for *de novo* atomic modelling revealed that the ordered cores of these filaments consist of residues 120-254 from the C-terminal, lumenal domain of TMEM106B and that the filaments are polymorphic. We solved the structures of TMEM106B filaments from the brains of 22 individuals with a neurodegenerative condition, and from the brains of two neurologically normal individuals (cases 1-24; **Methods; Table 1; Extended Data Table 1**). The neurodegenerative conditions for which we solved structures of TMEM106B filaments included sporadic and inherited Alzheimer’s disease (AD), pathological aging (PA), corticobasal degeneration (CBD), sporadic and inherited frontotemporal lobar degeneration (FTLD-TDP-A, FTLD-TDP-C and FTDP-17T), argyrophilic grain disease (AGD), limbic-predominant neuronal inclusion body 4R tauopathy (LNT), aging-related tau astrogliopathy (ARTAG), sporadic and inherited Parkinson’s disease (PD), dementia with Lewy bodies (DLB), multiple system atrophy (MSA), and amyotrophic lateral sclerosis (ALS). We observed three different TMEM106B protofilament folds (folds I-III; **Figure 1; Extended Data Figures 1–4**). Filaments with fold I are more common than filaments with fold II or fold III. For all three folds, we determined the structures of filaments that are made of a single protofilament. We also determined the structure of filaments comprising two protofilaments of fold I, related by C2 symmetry. In each individual, we only observed filaments with a single fold, without a clear relationship between folds and diseases.

**Table 1:** Overview of filament types identified in each case.

| Case | Disease | Age (yrs) | TMEM106B Filaments | Other Filaments |
| --- | --- | --- | --- | --- |
| 1 | AD | 79 | I-s (21%) / I-d (5%) | Aβ (37%) / Tau (37%) |
| 2 | FAD | 67 | I-s (16%) / I-d (1%) | Aβ (27%) / Tau (56%) |
| 3 | EOAD | 58 | I-s (31%) / I-d (<1%) | Aβ (15%) / Tau (54%) |
| 4 | PA | 59 | I-s (23%) / I-d (1%) | Aβ (76%) |
| 5 | CBD | 74 | I-s (6%) / I-d (1%) | Tau (93%) |
| 6 | CBD | 79 | I-s (11%) / I-d (1%) | Tau (88%) |
| 7 | FTDP-17T | 55 | I-s (23%) / I-d (3%) | Tau (74%) |
| 8 | AGD | 85 | III-s (8%) | Tau (92%) |
| 9 | AGD | 90 | I-s (29%) / I-d (3%) | Tau (68%) |
| 10 | LNT | 66 | I-s (17%) / I-d (2%) | Tau (81%) |
| 11 | ARTAG | 85 | III-s (67%) | Tau (11%) / Aβ (22%) |
| 12 | PD | 87 | III-s (13%) | Unknown (45%) / Tau (42%) |
| 13 | PDD | 64 | I-s (50%) / I-d (6%) | Aβ (28%) / Unknown (16%) |
| 14 | FPD | 67 | III-s (4%) | Unknown (96%) |
| 15 | DLB | 74 | III-s (36%) | Unknown (64%) |
| 16 | DLB | 73 | I-s (30%) / I-d (1%) | Aβ (62%) / Unknown (7%) |
| 17 | MSA | 85 | III-s (27%) | αS (73%) |
| 18 | MSA | 70 | I-s (13%) / I-d (5%) | αS (82%) |
| 19 | MSA | 68 | IIa-s (6%) / IIb-s (9%) / II-d (<1%) | αS (85%) |
| 20 | FTLD-TDP-A | 66 | I-s (21%) / I-d (30%) | Aβ (46%) / Unknown (3%) |
| 21 | FTLD-TDP-C | 65 | III-s (77%) | Unknown (23%) |
| 22 | ALS | 63 | III-s (46%) / III-d (10%) | Unknown (24%) / Aβ (19%) |
| 23 | Control | 75 | I-s (83%) / I-d (17%) | Undefined (<1%) |
| 24 | Control | 101 | I-s (92%) / I-d (8%) | Undefined (<1%) |

AD: sporadic Alzheimer's disease; FAD: familial Alzheimer's disease; EOAD: sporadic early-onset Alzheimer's disease; PA: pathological aging; CBD: corticobasal degeneration; FTDP-17T: familial frontotemporal dementia and parkinsonism linked to chromosome 17 caused by *MAPT* mutations; AGD: argyrophilic grain disease; LNT: limbic-predominant neuronal inclusion body 4R tauopathy; ARTAG: aging-related tau astrogliopathy; PD: sporadic Parkinson's disease; PDD: sporadic Parkinson's disease dementia; FPD: familial Parkinson's disease; DLB: dementia with Lewy bodies; MSA: multiple system atrophy; FTLD-TDP-A: familial frontotemporal lobar degeneration with TDP-43 inclusions type A; FTLD-TDP-C: sporadic frontotemporal lobar degeneration with TDP-43 inclusions type C; ALS: amyotrophic lateral sclerosis; Control: neurologically normal individual; $\alpha$ S: α-synuclein. TMEM106B filaments are indicated according to their protofilament fold (I-III) and whether they comprise one (-s) or two (-d) protofilaments. Percentages of protein filament types were calculated based on the number of extracted segments from manually picked filaments (and in some cases on the number of segments after 2D classification to separate TMEM106B filaments comprising one or two protofilaments). These values may not reflect what is present in the brain, nor be directly comparable between cases.

**Figure 1:**
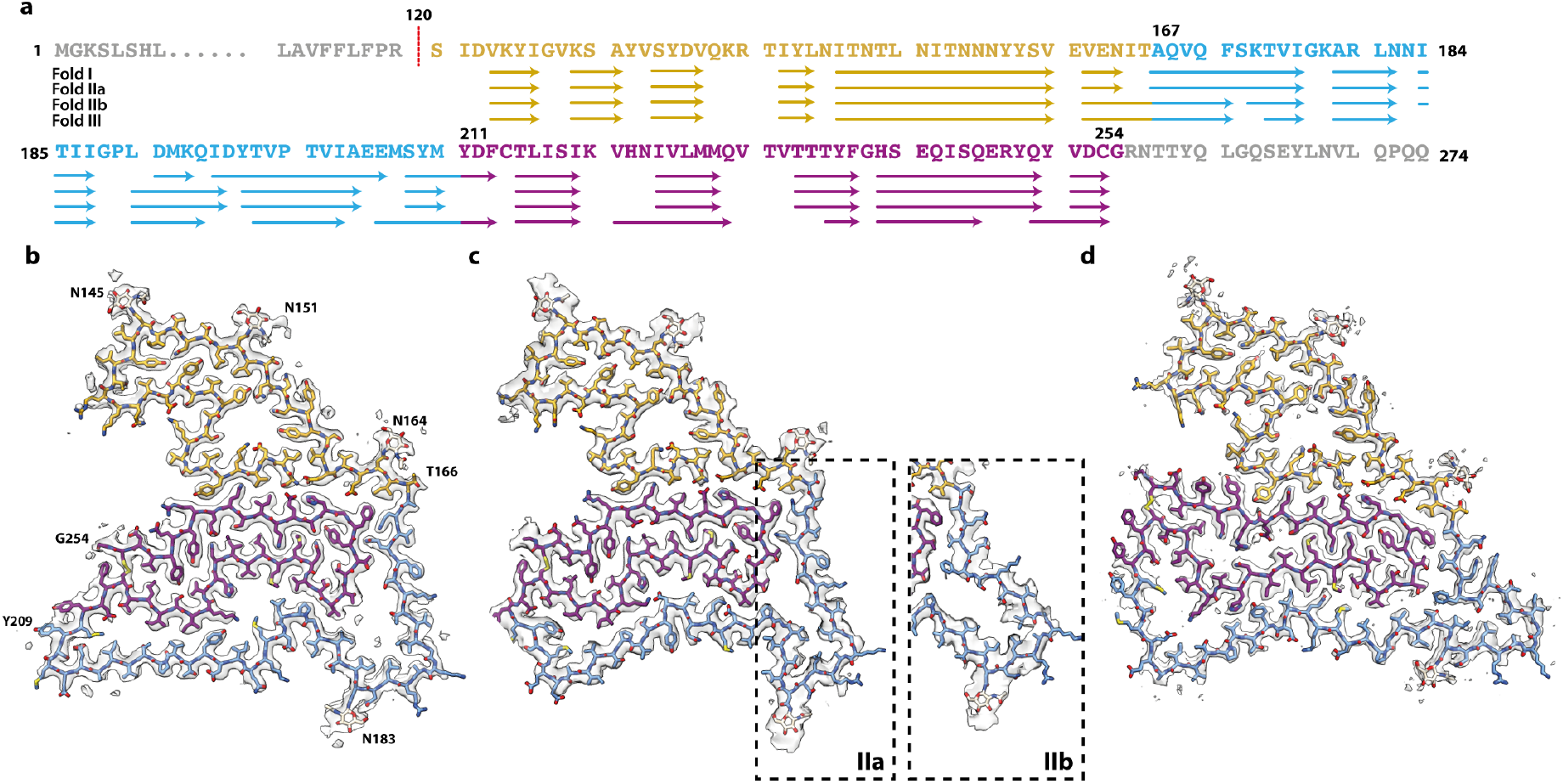
Three TMEM106B protofilament folds. **a.** Amino acid sequence of TMEM106B, with residues that form β-strands in the I, IIa, IIb and III folds indicated with arrows. **b-d.** Cryo-EM density map (in transparent grey) and atomic models for TMEM106B protofilament folds I (b), II (c) and III (d). Two alternative conformations of fold II (IIa and IIb) are indicated within dashed boxes. Residues 120-166 are shown in yellow; residues 167-210 in light blue and residues 211-254 in magenta.

The TMEM106B folds share a similar five-layered ordered core comprising residues S120-G254 and contain 17 β-strands, each ranging between 3 and 15 residues. Our best maps for filaments with folds I, II and III had resolutions of 2.6, 3.4 and 2.8 Å, and came from case 1 (sporadic AD), case 19 (MSA) and case 17 (MSA), respectively. TMEM106B remains fully glycosylated in all folds, as there are large additional densities corresponding to glycan chains attached to the side chains of asparagines 145, 151, 164 and 183. The fifth glycosylation site at N256 is outside the ordered core, with the C-terminal twenty residues being probably disordered. We divide the sequence that forms the ordered cores of the folds into three regions according to their degree of structural conservation: the N-terminal region (S120-T166) is conserved in all three folds; the C-terminal region (Y211-G254) is only conserved in folds I and II; and the middle region (A167-M210) varies between folds.

The N-terminal region, S120-T166, forms the first two layers of the five-layered ordered cores. It comprises five short and one longer β-strands that constitute a tightly packed core with hydrophobic and neutral polar residues on one side, and a large polar cavity that is filled by solvent on the other side. The three glycosylation sites in this region are located in the outer layer, adopting an extended conformation. N-terminal residue S120 in the inner layer is buried inside the ordered core, where it packs closely against E161 from the N-terminal region and H239 and E241 from the C-terminal region (**Extended Data Figure 5)**.

The C-terminal region, Y211-G254, forms the two central layers of the ordered cores. It adopts a compact hairpin-like structure, the ends of which are held together by a disulfide bond between C214 and C253. Segment F237-E246 that packs against the N-terminal region has the same conformation in all three folds, whereas in the rest of the hairpin-like structure, 15 residues have opposite ‘inward/outward’ orientations in fold III compared to folds I and II. Moreover, despite similar interfaces between N- and C-terminal parts in all three folds, these regions in fold III are separated along the filament axis by one more rung than in folds I and II (**Extended Data Figure 3e**).

The middle region, A167-M210, forms the fifth layer of the ordered cores and contains the fourth glycosylation site at N183. In fold I, this region packs loosely against the other side of the C-terminal hairpin-like region with the formation of three large amphipathic cavities. In fold II, these internal cavities are smaller than in fold I. We observed two subtypes of fold II (IIa and IIb) that differ mainly by the conformation of segment A167-I187. The packing of the middle region against the C-terminal region is tightest in fold III, leaving only one sizable cavity with a salt bridge between E206 and K220. Only fold III shows cis-isomerisation of P189. In folds IIb and III, there is a large extra density at the end of the side chain of K178, suggesting that this residue may be post-translationally modified. Likewise, there is an additional density in front of the side chain of Y209 in fold I, but not in the other folds (**Extended Data Figure 1**). It is possible that these residues determine the formation of the different folds.

In all three folds, residues E177-N183 adopt a conserved conformation, with positively charged residues K178 and R180 pointing outwards. In filaments made of two protofilaments with fold I, two pairs of these residues are on opposite sides of a contiguous additional density that runs along the helical symmetry axis. Because the cofactor responsible for this density probably does not obey the helical symmetry imposed, the map in this region is of insufficient quality to allow its identification. Nevertheless, the density’s shape and presumed charge are consistent with the presence of a sulfated glycosaminoglycan. Although we did not solve the structures of filaments comprising two protofilaments of folds II or III, the micrographs of case 19, the only case for which we observed filaments with fold IIa/b, and the micrographs of case 21, FTLD-TDP-C with fold III, also contained wider filaments that probably comprised two TMEM106B protofilaments (**Extended Data Figure 6**).

In the absence of an experimentally determined native structure for TMEM106B, we examined its structure as predicted by AlphaFold (Tunyasuvunakool et al., 2021) (**Extended Data Figure 7**). Whereas the formation of amyloid filaments in the brain is often associated with natively unfolded proteins or low-complexity domains, the sequence S120-G254, which spans the ordered core of TMEM106B filaments, is confidently predicted to be a globular domain of the immunoglobulin-like β-sandwich fold. Glycosylation sites at N145, N151, N164 and N183 are positioned on the outside of the fold, and the disulfide bond between C214 and C253 is also predicted to form in the native structure. The β-sandwich domain is connected to a single transmembrane helix, without a flexible linker sequence. Moreover, there is a hydrophobic surface patch at this end of the domain, suggesting that it is positioned close to the membrane. It thus seems unlikely that the cleavage site at S120, the buried N-terminal residue in all TMEM106B filaments, can be accessed by lysosomal proteases. Shedding of the lumenal domain may therefore happen in a non-canonical way.

We previously showed that distinct amyloid folds of tau, α-synuclein, and Aβ characterise different neurodegenerative diseases (Fitzpatrick et al., 2017; Falcon et al., 2018; Falcon et al., 2019; Zhang et al., 2020; Schweighauser et al., 2020; Shi et al., 2021; Yang et al., 2021). We now describe the presence of TMEM106B filaments in many of these diseases, without a correlation between folds and diseases. Therefore, we also examined 13 brains from neurologically normal individuals, varying in age from 20 to 101 years. By immunoblotting the sarkosyl-insoluble fraction for the diseased cases with an antibody specific for residues 239-250 of TMEM106B, we observed a band of 29 kDa, which probably corresponds to the 17 kDa C-terminal fragment plus 12 kDa of glycosylation and other modifications (**Figure 2; Extended Data Figure 8**). This band was not present in the brains from neurologically normal individuals who were less than 46 years of age, rendering unlikely the possibility that TMEM106B aggregation was an artefact caused by tissue extraction. In future, it will be important to demonstrate the presence of TMEM106B filaments in brain tissue. We consistently observed the 29 kDa band in the brains of control individuals older than 69 years. Cryo-EM structure determination for two of these cases, aged 75 and 101 years, confirmed the presence of TMEM106B filaments, comprising one or two protofilaments with fold I. Interestingly, the 29 kDa band was not present in the frontal cortex from a 15-year-old individual with early-onset dementia with Lewy bodies (Takao et al., 2004).

**Figure 2:**
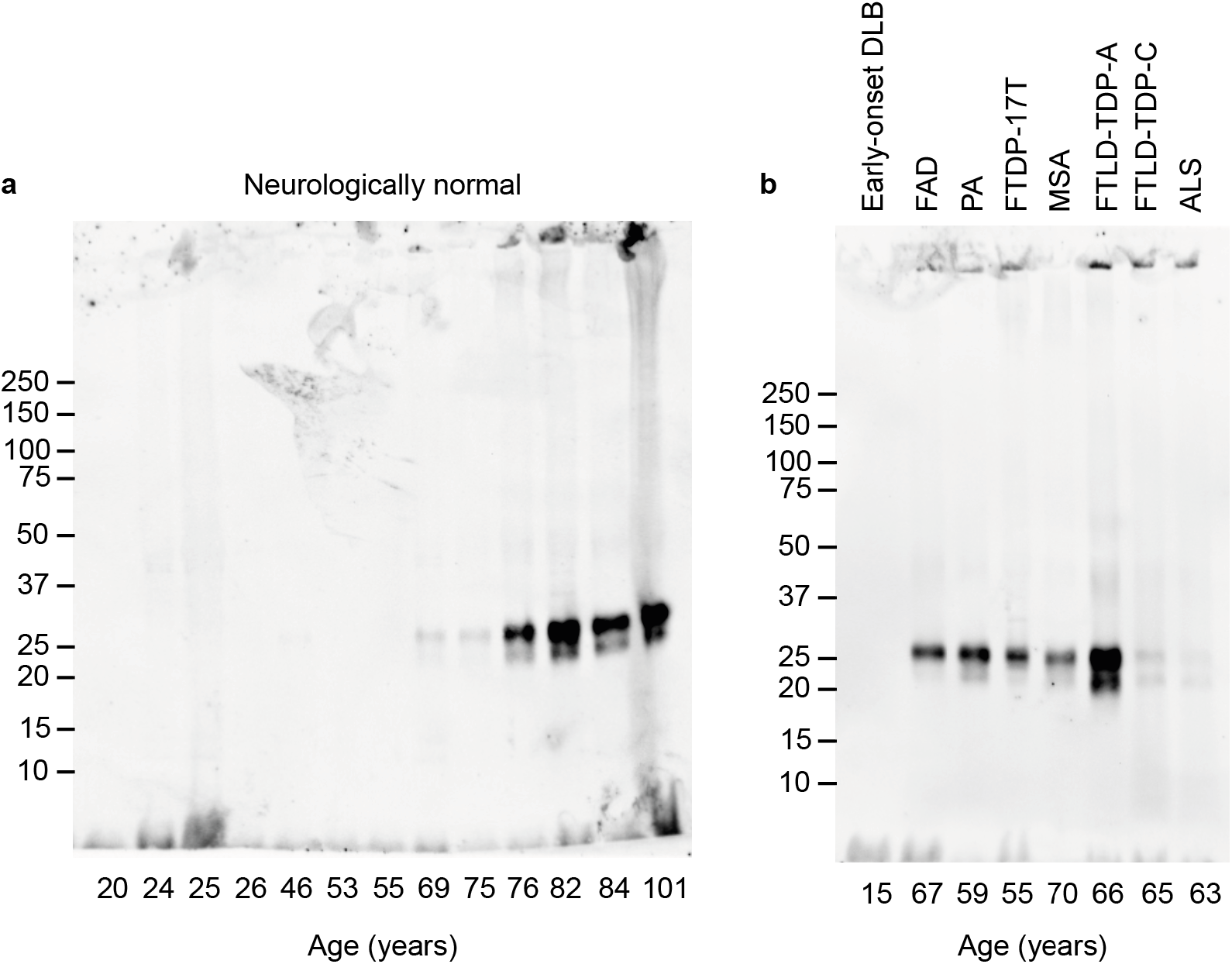
TMEM106B Western blots. **a.** Immunoblot analysis with anti-TMEM239 (residues 239-250) of sarkosyl-insoluble extract from brains of 13 neurologically normal individuals aged 20-101 years. Cryo-EM structures were determined from individuals aged 75 and 101. **b.** Immunoblot analysis of sarkosyl-insoluble extracts from 8 individuals with abundant amyloid deposits (cases: early onset DLB, FAD, PA, FTDP-17T, MSA, FTLD-TDP-A, FTLD-TDP-C, ALS).

Our results suggest that amyloid filaments of the lysosomal protein TMEM106B form in an age-dependent manner in human brain, without a clear mechanistic connection to disease. Until now, the presence of abundant amyloid filaments in human tissues has typically been associated with disease. Mutations in the genes for tau, APP, α-synuclein and TDP-43, as well as in the genes encoding enzymes, like the presenilins, that process APP, are causative of neurodegenerative diseases. In addition, cryo-EM structures of amyloid filaments made of these proteins display distinct folds that are characteristic of different diseases. Although TMEM106B has been associated with frontotemporal dementia and other diseases, the evidence for a causal relationship between TMEM106B aggregation and disease remains unclear, and distinct TMEM106B folds do not characterise different diseases. Instead, our observations suggest that TMEM106B filaments form in an age-dependent manner. Like lipofuscin, a lysosomal complex of oxidised proteins and lipids that develops in an age-dependent manner in many tissues (Jung et al., 2007), TMEM106B filaments may also form in lysosomes. Lysosomal dysfunction has been implicated in the pathogenesis of neurodegenerative diseases (Nixon, 2013). Additional studies are needed to determine if TMEM106B filaments can be found in tissues other than the brain and to assess the role of filament formation in relation to human aging and pathologies.

## Acknowledgements

We thank the patients’ families for donating brain tissues; U. Kuederli, M. Jacobsen, F. Epperson and R.M. Richardson for human brain collection and technical support; E. Gelpi for preparing brain samples from the ARTAG case; T. Darling and J. Grimmett for help with high-performance computing; I. Lavenir, J. Macdonald, M. Bacioglu, M.G. Spillantini, and R. A. Crowther for helpful discussions; the EM facility of the Medical Research Council (MRC) Laboratory of Molecular Biology for help with cryo-EM data acquisition; the Edinburgh Brain and Tissue Bank, which is supported by the MRC, for providing brain samples of neurologically normal controls. We acknowledge Diamond Light Source for access and support of the cryo-EM facilities at the UK’s national Electron Bio-imaging Centre (eBIC) [under proposals EM17434-75 and BI23268-49], funded by Wellcome Trust, MRC and BBSRC. This work is supported by the MRC (MC_UP_120/25 to B.F., MC_U105184291 to M.G. and MC_UP_A025_1013 to S.H.W.S.), and the EU/EFPIA/Innovative Medicines Initiative [2] Joint Undertaking IMPRiND (project 116060), to M.G., the Japan Agency for Science and Technology (Crest, JPMJCR18H3), to M.H., the Japan Agency for Medical Research and Development (AMED, JP20dm0207072), to M.H., and (AMED, JP21wm0425019), to M.T., the Japan Society for the Promotion of Science (JSPS, Kakenhi 21K06417), to M.T., the U.S. National Institutes of Health (P30-AG010133, UO1-NS110437 and RF1-AG071177), to R.V. and B.G., and the Department of Pathology and Laboratory Medicine, Indiana University School of Medicine, to R.V. and K.L.N. and B.G.. G.G.K. was supported by the Safra Foundation and the Rossy Foundation. T.R. is supported by the National Institute for Health Research Queen Square Biomedical Research Unit in Dementia. M.T. is supported by intramural funds from the National Center of Neurology and Psychiatry. T.L. holds an Alzheimer’s Research UK Senior Fellowship. The Queen Square Brain Bank is supported by the Reta Lila Weston Institute for Neurological Studies.

For the purpose of open access, the authors have applied a CC-BY public copyright license to any Author Accepted Manuscript version arising.

## Author contributions

K.L.N., S.M., M.M., Y.Saito, M.Y.,K.H., T.L., T.R., G.G.K., J.v.S., M.T., M.H., B.G., B.F. and M.G. identified patients and performed neuropathology; M.S., Y.Shi, M.H., D.A., Y.Y., W.Z., H.J.G., R.V., G.I.H., A.T. and M.H. performed brain sample analysis; M.S., Y.Shi, D.A., Y.Y., W.Z. and A.K. collected cryo-EM data; M.S., Y.Shi, D.A., Y.Y., S.L., W.Z., B.F., A.G.M. and S.H.W.S. analysed cryo-EM data; M.S. and M.H. performed immunoblot analysis; S.L. performed recombinant protein expression and epitope mapping; M.G. and S.H.W.S. supervised the project. All authors contributed to the writing of the manuscript.

## Methods

### Clinical history and neuropathology

We determined the cryo-EM structures of TMEM106B filaments from the brains of 24 individuals (**Table 1, Extended Data Table 1**). Most individuals have been reported (Falcon et al., 2018; Zhang et al., 2020; Schweighauser et al., 2020; Shi et al., 2021; Yang et al., 2021). Unpublished cases are described below. Early-onset Alzheimer’s disease (EOAD; case 3) was in a 58-year-old woman who died with a neuropathologically confirmed diagnosis following a 7-year history of memory loss. FTDP-17T (case 7) was in a 55-year-old man who died with a neuropathologically confirmed diagnosis following a 2-year history of behavioural changes, aphasia and dementia caused by a P301L mutation in *MAPT*. His brother, sister and mother were also affected. Sporadic Parkinson’s disease (case 12) was in a 87-year-old male who died with a neuropathologically confirmed diagnosis following an 8-year history of Parkinson’s disease. Inherited Parkinson’s disease (case 14) was in a 67-year-old woman who died with a neuropathologically confirmed diagnosis following a 10-year history of Parkinson’s disease caused by a G51D mutation in *SNCA*. FTLD-TDP-C (case 21) was in a 65-year-old woman who died with a neuropathologically confirmed diagnosis following a 9-year history of semantic dementia. ALS (case 22) was in a 63-year-old woman who died with a neuropathologically confirmed diagnosis of ALS stage 4, type B TDP-43 pathology, following a 2-year and 5-month history of motor symptoms, without dementia. Control 1 (case 23) was a 75-year-old man who died of coronary heart disease without significant neuropathological abnormalities. Control 2 (case 24) was a 101-year-old man with mild tau pathology (Braak stage 1) and mild cerebral amyloid angiopathy.

### Extraction of TMEM106B filaments

Sarkosyl-insoluble material was extracted from frontal cortex (EOAD, FTLD-TDP-C, control cases 1 and 2), cingulate cortex (sporadic PD), temporal cortex (inherited PD, FTDP-17T) and motor cortex (ALS), essentially as described (Tarutani et al., 2018). Similar extraction methods were used for all other cases, which have been described in the references in Extended Data Table 1. The original sarkosyl extraction method, which we used in our work on the cryo-EM structures of tau filaments from Alzheimer’s disease, chronic traumatic encephalopathy and Pick’s disease (Fitzpatrick et al., 2017; Falcon et al., 2018; Falcon et al., 2019), uses sarkosyl only after the first, low-speed centrifugation step (Greenberg and Davies, 1990). The method of Tarutani et al. also uses sarkosyl at the beginning, i.e. before the first centrifugation step. This protocol change was essential for detecting abundant TMEM106B filaments. In brief, tissues were homogenised in 20 volumes (v/w) extraction buffer consisting of 10 mM Tris-HCl, pH 7.5, 0.8 M NaCl, 10% sucrose and 1 mM EGTA. Homogenates were brought to 2% sarkosyl and incubated for 30 min. at 37° C. Following a 10 min. centrifugation at 10,000 g, the supernatants were spun at 100,000 g for 20 min. The pellets were resuspended in 700 μl/g extraction buffer and centrifuged at 5,000 g for 5 min. The supernatants were diluted 3-fold in 50 mM Tris-HCl, pH 7.5, containing 0.15 M NaCl, 10% sucrose and 0.2% sarkosyl, and spun at 166,000 g for 30 min. Sarkosyl-insoluble pellets were resuspended in 50 μl/g of 20 mM Tris-HCl, pH 7.4 containing 100 mM NaCl.

### Immunoblotting

Immunoblotting was carried out as described (Tanugushi-Watanabe et al., 2016). Sarkosyl-insoluble pellets were sonicated for 5 min at 50% amplitude (QSonica). They were resolved on 4-20% Tris-glycine gels (Novex) and antibody TMEM239 (a rabbit polyclonal antibody that was raised against a synthetic peptide corresponding to amino acids 239-250 of TMEM106B) was used at 1:2,000.

### Cloning

TMEM106B C-terminal fragment (120 – 274) incorporated in pET3A was purchased from Genscript™. The corresponding epitope-deletion construct, lacking residues 239-250 (Δ239-250), was made using an *in vivo* assembly approach (García-Nafría et al., 2016). Forward and reverse primers were obtained from Integrated DNA Technologies and were designed to share 15-20 nucleotides of homologous region and 15-30 nucleotides for annealing to the template, flanking the region of deletion, with melting temperatures ranging from 58 to 65 °C. Prior to transformation, PCR products were treated with *DpnI*.

### Purification of recombinant TMEM106B

Plasmids were transformed into *E. coli* BL21 (DE3pLys) (Agilent). One plate was used to inoculate 500 ml terrific broth (TB), supplemented with 2.5 mM MgSO_4_ and 2 % ethanol, 100 mg/l ampicillin, and grown with shaking at 220 rpm at 37 °C, until an OD of 0.8 was reached; expression was induced with 1 mM IPTG for 4 h at 37 °C. Cells were harvested by centrifugation for 20 min at 4,000 x g at 4 °C and resuspended in cold buffer A: 4X PBS, pH 7.4, 25 mM DTT, 0.1 mM PMSF and cOmplete protease inhibitor tablets (4 tablets per 100 ml). Resuspension was performed using a Polytron with a 10:1 volume-to-weight ratio of pellet to buffer. The homogenised pellets were sonicated (40 % amplitude, 5 s on, 10 s off, for 6 min) at 4 °C. Lysed cells were then centrifuged at 30,000 x g for 40 min at 4 °C, and the pellets were resuspended in buffer A plus 2 M urea and 2 % Triton, incubated for 30 min at 40 °C and centrifuged at 30,000 x g for 20 min at 25 °C. This resuspension step was repeated three times. Subsequently, the pellets, appearing as dense white matter indicative of inclusion bodies, were resuspended in buffer A plus 2 M urea, incubated for 30 min at 40 °C and centrifuged at 30,000 x g for 20 min at 25 °C. Finally, the pellets were resuspended in buffer A plus 8 M urea, using a 20:1 volume-to-weight ratio, for 1 h with shaking at 100 rpm at 60 °C, and centrifuged at 30,000 x g for 20 min at 25 °C. These pellets were resuspended in 4X PBS, pH 7.4, 50 mM DTT and 8 M urea, and left shaking overnight at 100 rpm at 60 °C. The resuspended pellets were centrifuged at 45,000 x g for 30 min and the supernatants concentrated, followed by buffer-exchange using a PD10 desalting column into 2X PBS, pH 7.4 and 50 mM DTT. Samples were further concentrated to 3 mg/ml using a 3kDa cutoff molecular weight concentrator, and used for Western blots to confirm the specificity of the generated antibody TMEM239 (**Extended Data Figure 8**).

### Electron cryo-microscopy

For all cases, except EOAD, FTDP-17T, LNT, sporadic PD, inherited PD, FTLD-TDP-C, ALS and control cases 1 and 2, the cryo-EM data sets have been described in the references in **Extended Data Table 1**. For the remaining cases, resuspended sarkosyl insoluble pellets were applied to glow-discharged holey carbon gold grids (Quantifoil R1.2/1.3, 300 mesh) and plunge-frozen in liquid ethane using an FEI Vitrobot Mark IV. FTLD-TDP-A, FTLD-TDP-C, and ALS samples were treated with 0.4 mg/ml pronase for 50-60 min prior to glow-discharging, which further improved the TMEM106B filament yield. Images for cases EOAD, FTDP-17T, LNT, FPD, FTLD-TDP-C and ALS were acquired on Thermo Fisher Titan Krios microscopes, operated at 300 kV, with a Gatan K2 or K3 detector in counting mode, using a Quantum energy filter (Gatan) with a slit width of 20 eV to remove inelastically scattered electrons. Images for EOAD, sporadic PD and control cases 1 and 2 were acquired on a Thermo Fisher Titan Krios, operated at 300 kV, using a Falcon-4 detector and no energy filter.

### Helical reconstruction

Movie frames were gain-corrected, aligned, dose-weighted and then summed into a single micrograph using RELION’s own motion correction program (Zivanov et al., 2019). The micrographs were used to estimate the contrast transfer function (CTF) using CTFFIND-4.1 (Rohou and Grigorieff, 2015). All subsequent image-processing steps were performed using helical reconstruction methods in RELION (He and Scheres, 2017, Zivanov et al., 2018). TMEM106B filaments were picked manually, as they could be distinguished from filaments made of other proteins by their general appearance and the apparent lack of a fuzzy coat. TMEM106B filaments comprising one or two protofilaments were picked separately. For all data sets, reference-free 2D classification was performed to select suitable segments for further processing. Initial 3D reference models were generated *de novo* from the 2D class averages using an estimated rise of 4.75 Å and helical twists according to the observed cross-over distances of the filaments in the micrographs (Scheres, 2020) for datasets case 10 (LNT; folds I-s and I-d), case 18 (MSA; fold I-d), case 19 (MSA; folds IIa and IIb), case 17 (MSA; fold III). Refined models from these cases, low-pass filtered to 10-20 Å, were used as initial models for the remaining cases. Combinations of 3D auto-refinements and 3D classifications were used to select the best segments for each structure. For all data sets, Bayesian polishing (Zivanov et al., 2019) and CTF refinement (Zivanov et al., 2020) were performed to further increase the resolution of the reconstructions. Final reconstructions were sharpened using the standard post-processing procedures in RELION, and overall final resolutions were estimated from Fourier shell correlations at 0.143 between the two independently refined halfmaps, using phase-randomisation to correct for convolution effects of a generous, soft-edged solvent mask (Chen et al., 2013). Further details of data acquisition and processing for the data sets that resulted in the best maps for five different TMEM106B filaments (filaments made of one or two protofilaments with fold I, as well as filaments made of one protofilament with fold IIa, fold IIb or fold III) are given in **Extended Data Table 2**.

### Model building

TMEM106B was identified by scanning the human proteome with different sequence motifs (de Castro et al., 2006), deduced from initial maps of folds I and III. A simple combination of four N-glycosylation motifs N-x-[TS] with the exact spacers, N-x-[ST]-x(3)-N-x-[ST]-x(10)-N-x-[ST]-x(16)-N-x-[ST], was the most effective, resulting in only a single hit for TMEM106, the sequence of which corresponded well to the entire maps. Atomic models comprising three β-sheet rungs were built *de novo* in Coot (Casañal et al., 2020) in the best available map for each of the five different structures. Coordinate refinement was performed in ISOLDE (Croll, 2018). Dihedral angles from the middle rung, which was set as a template in ISOLDE, were also applied to the rungs below and above. For each refined structure, separate model refinements were performed for the first half map, after increasing the temperature to 300 K for 1 minute, and the resulting model was then compared to that same half map (FSC_work_) as well as the other half-map (FSC_test_) to confirm the absence of overfitting. Final statistics for the refined models are given in **Extended Data Table 2.**

### Ethical review processes and informed consent

Studies carried out at Indiana University, Tokyo Metropolitan Institute of Medical Science, Tokyo National Center Hospital, UCL Queen Square Institute of Neurology, Toronto University, Vienna Medical University, Rotterdam University, and the Edinburgh Brain and Tissue Bank were approved through the ethical review processes at each Institution. Informed consent was obtained from the patients’ next of kin.

### Data availability

Cryo-EM maps have been deposited in the Electron Microscopy Data Bank (EMDB) under accession numbers XXX. Corresponding refined atomic models have been deposited in the Protein Data Bank (PDB) under accession numbers XXX.

## Extended Data Figures

**Extended Data Figure 1:**
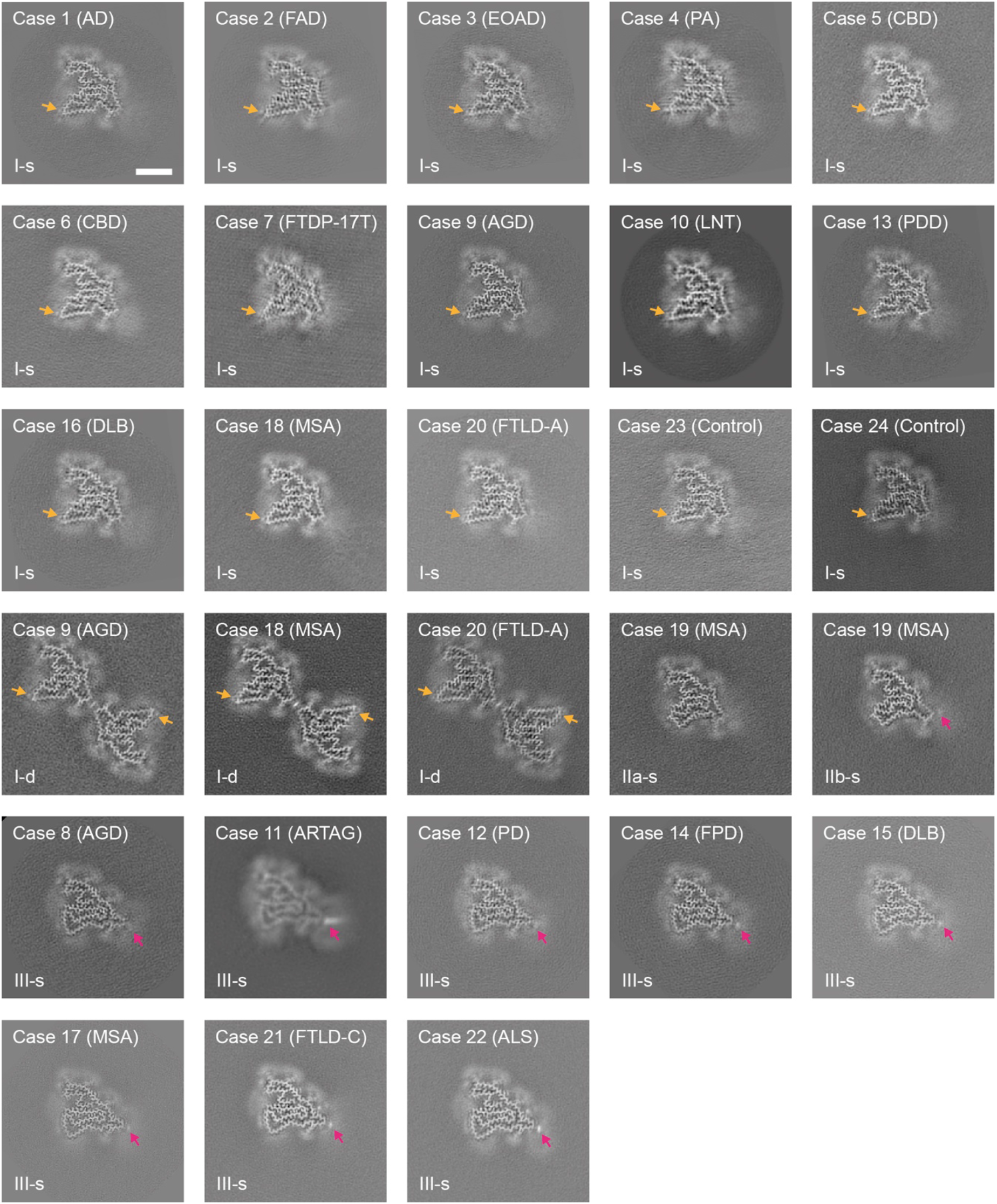
TMEM106B reconstructions. Cross-sections of TMEM106B filament reconstructions, perpendicular to the helical axis and with a projected thickness of approximately one β-rung, for all cases examined. Orange arrows point at additional densities in front of Y209; magenta arrows point at additional densities in front of K178. Scale bar 5nm.

**Extended Data Figure 2:**
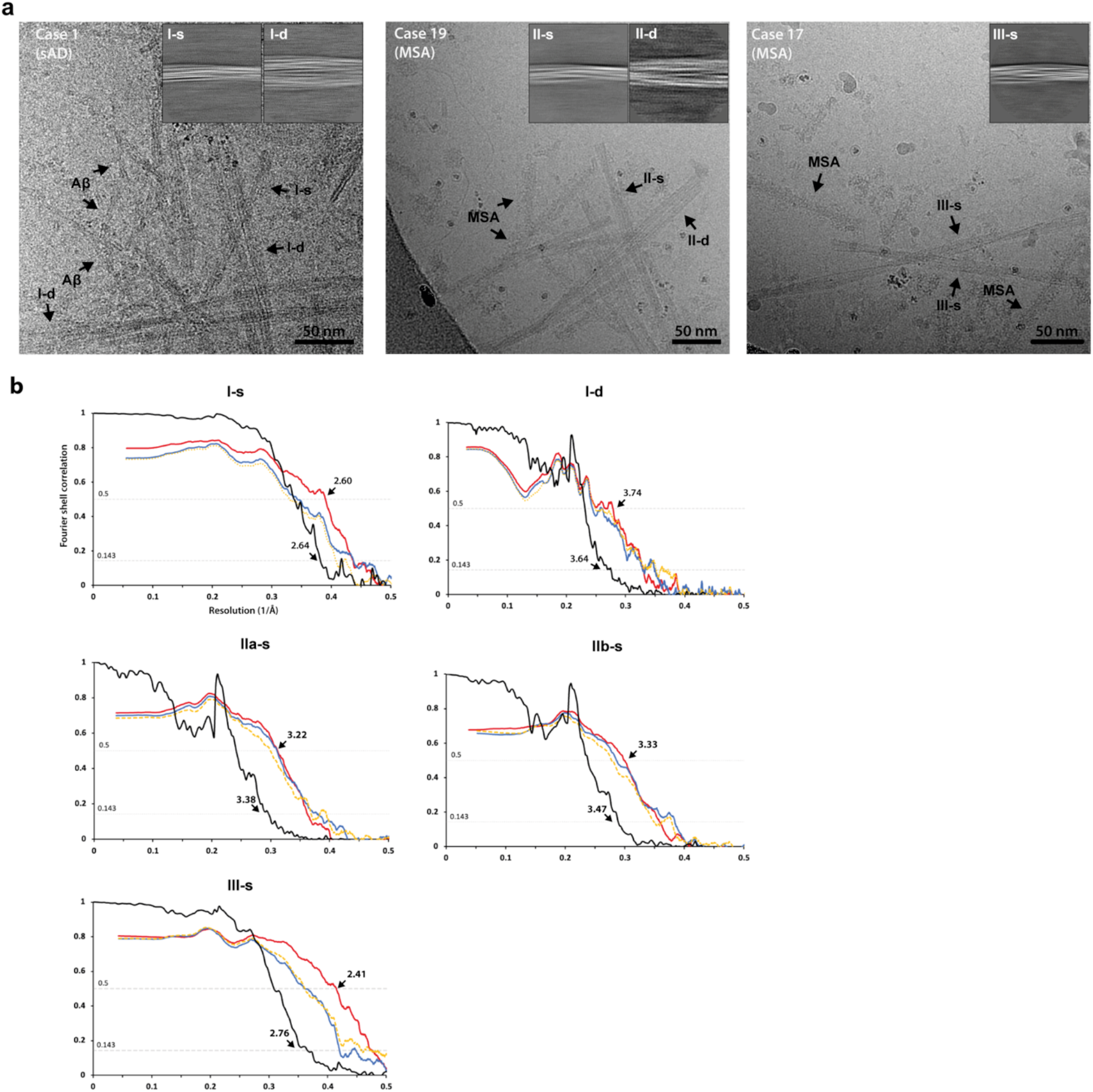
Cryo-EM images and resolution estimates. **a.** Cryo-EM micrographs of cases 1, 19 and 17, with insets showing representative 2D class averages of TMEM106B filaments I-s, I-d, II-s, II-d and III-s. Examples of the different types of TMEM106B filaments, as well as filaments of Aβ and MSA filaments of α-synuclein, are indicated in the micrographs with black arrows. Scale bars 50nm. **b.** Fourier shell correlation (FSC) curves for cryo-EM maps and structures of TMEM106B filaments I-s, I-d, II-s, II-d and III-s. FSC curves for two independently refined cryo-EM half maps are shown in black; for the final refined atomic model against the final cryo-EM map in red; for the atomic model refined in the first half map against that half map in blue; and for the refined atomic model in the first half map against the other half map in yellow.

**Extended Data Figure 3:**
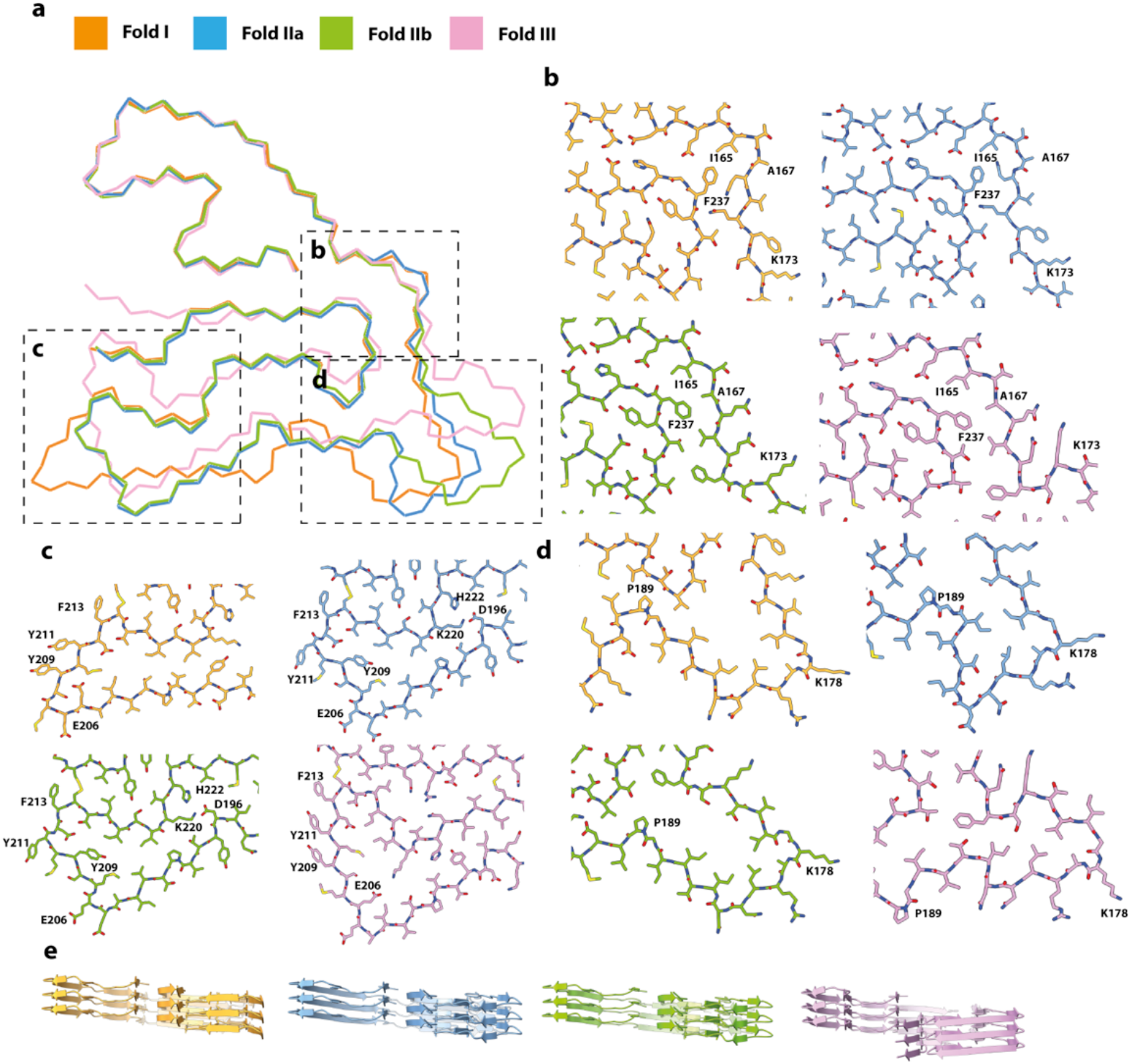
Comparisons of TMEM106B folds I-III. **a** Ribbon view of folds I-III aligned at residues 120-166 (centre) with close-up views for three regions (**b-d**). **e.** Cartoon views for three subsequent β-rungs for each fold, viewed from the left-hand side of the ribbon view in panel A. Fold I is shown in orange; fold IIa in blue; fold IIb in green; fold III in pink.

**Extended Data Figure 4:**
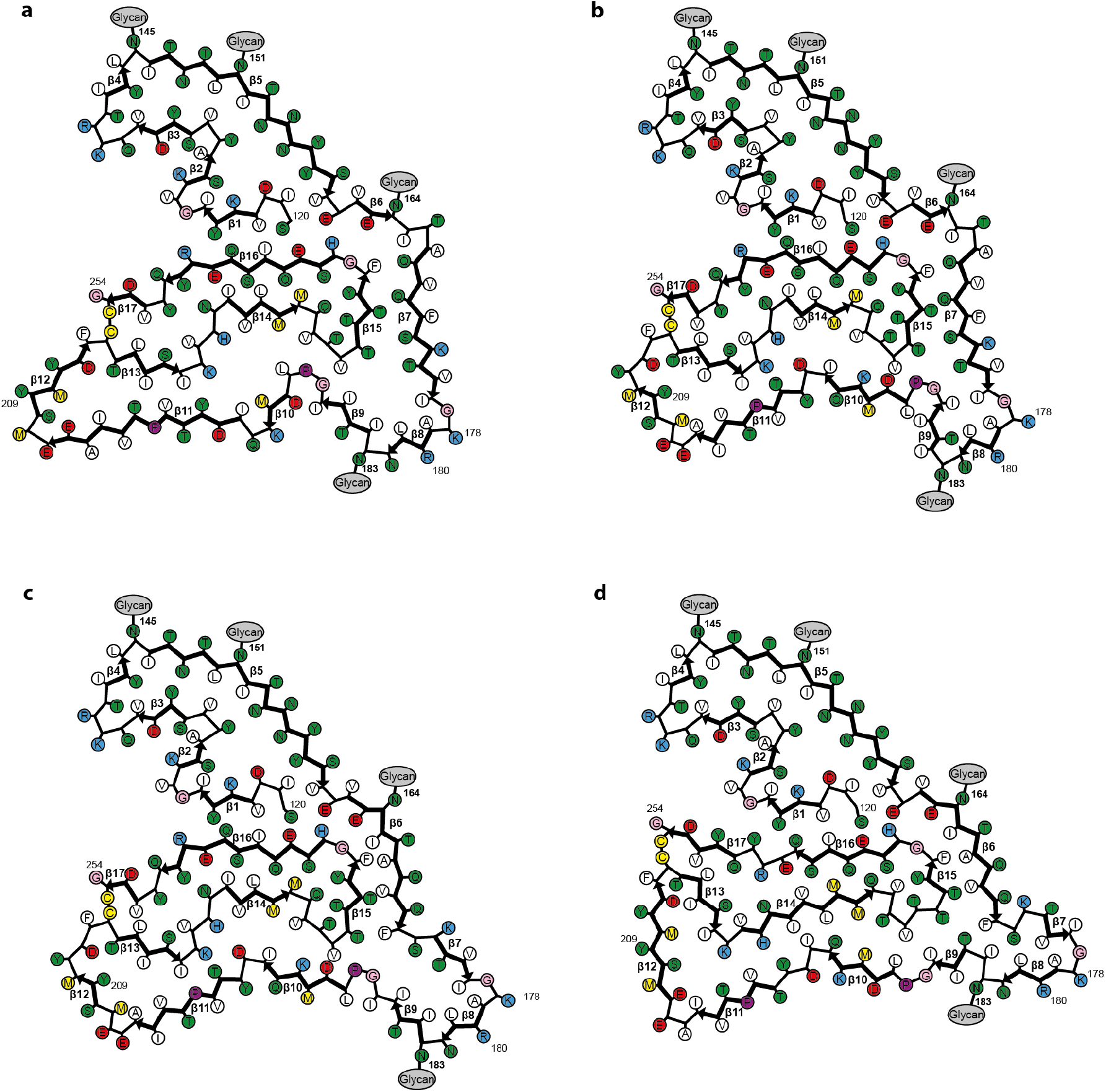
Schematics of the different folds. Schematics of fold I (**a**), fold IIa (**b**), fold IIb (**c**) and fold III (**d**). Negatively charged residues are shown in red, positively charged residues in blue, polar residues in green, apolar residues in white, sulfur-containing residues in yellow, prolines in purple, and glycines in pink. Thick connecting lines with arrowheads indicate β-strands.

**Extended Data Figure 5:**
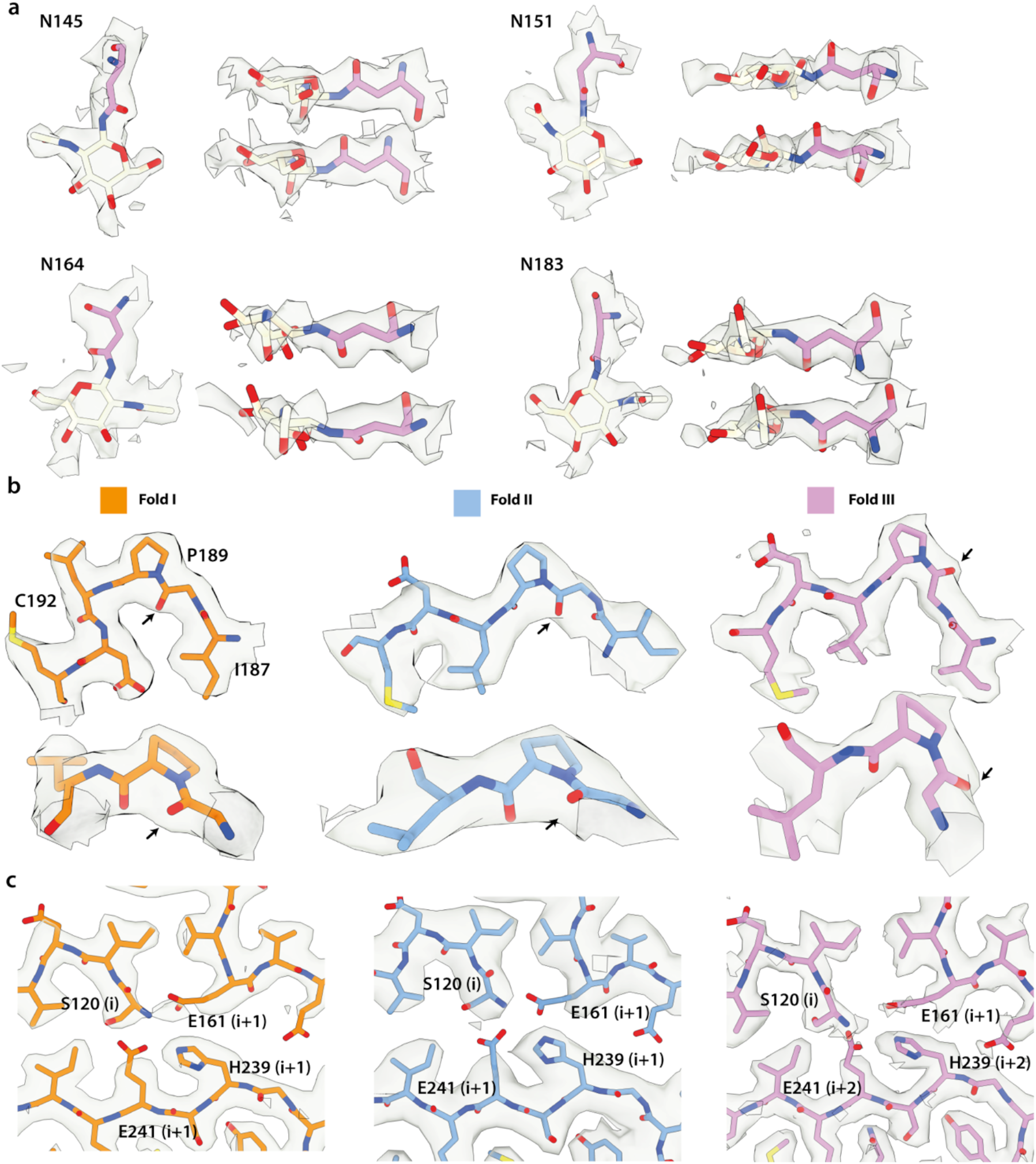
Close-up views of cryo-EM densities. **a.** Cryo-EM densities (transparent grey) for N-linked glycosylation of asparagines 145, 151, 164 and 183 in fold III, showing the first, most ordered glcNac saccharide of the glycan chains. **b.** Density for residues 187-192 in fold I (left, orange), fold II (middle, blue) and fold III (right, pink), as viewed from the top (top panels) and the side (bottom panels). Carbonyl oxygens of P189 are indicated with black arrows. In folds I and II, P189 adopts a *trans*-configuration, whereas in fold III, P189 adopts a *cis*-conformation. **c.** Density for the N-terminal S120 residue, and its surrounding residues E161, H239, E241. The latter are one β-rung above (i+1) the β-rung of S120 (i) in folds I and II. In fold III, H239 and E241 are two β-rungs above (i+2) the β-rung of S120.

**Extended Data Figure 6:**
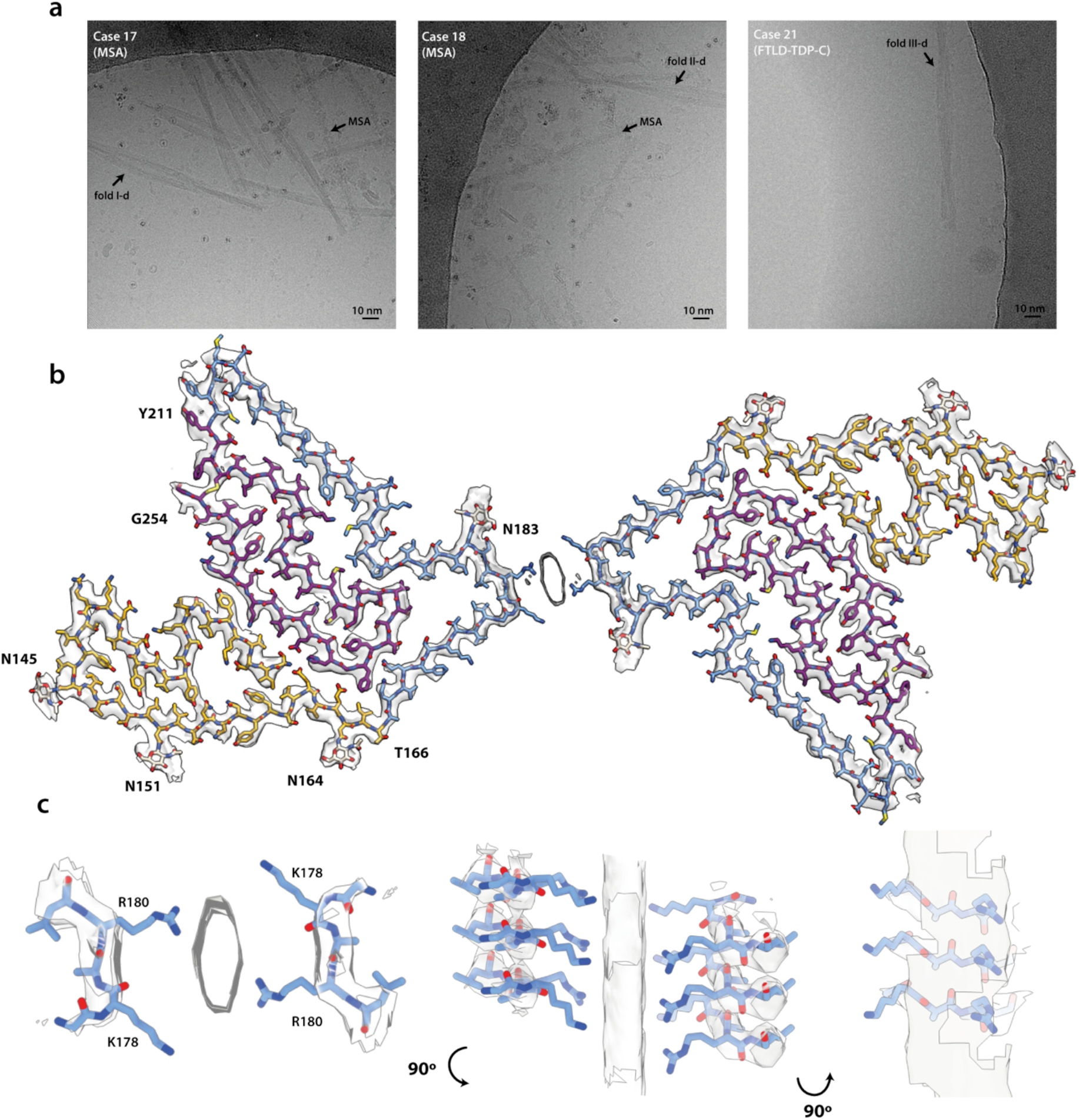
TMEM106B filaments comprising two protofilaments. **a.** Cryo-EM micrographs with filaments comprising two protofilaments, with fold I for MSA case 18 (I-d; left), with putative fold II for MSA case 19 (II-d; middle), and with putative fold III for XXX (III-d; right); α-synuclein filaments typical for MSA are also indicated (MSA). **b.** Cryo-EM density map and atomic model of TMEM106B filaments comprising two protofilaments of fold I. **c.** Three orthogional close-up views of the inter-protofilament interface.

**Extended Data Figure 7:**
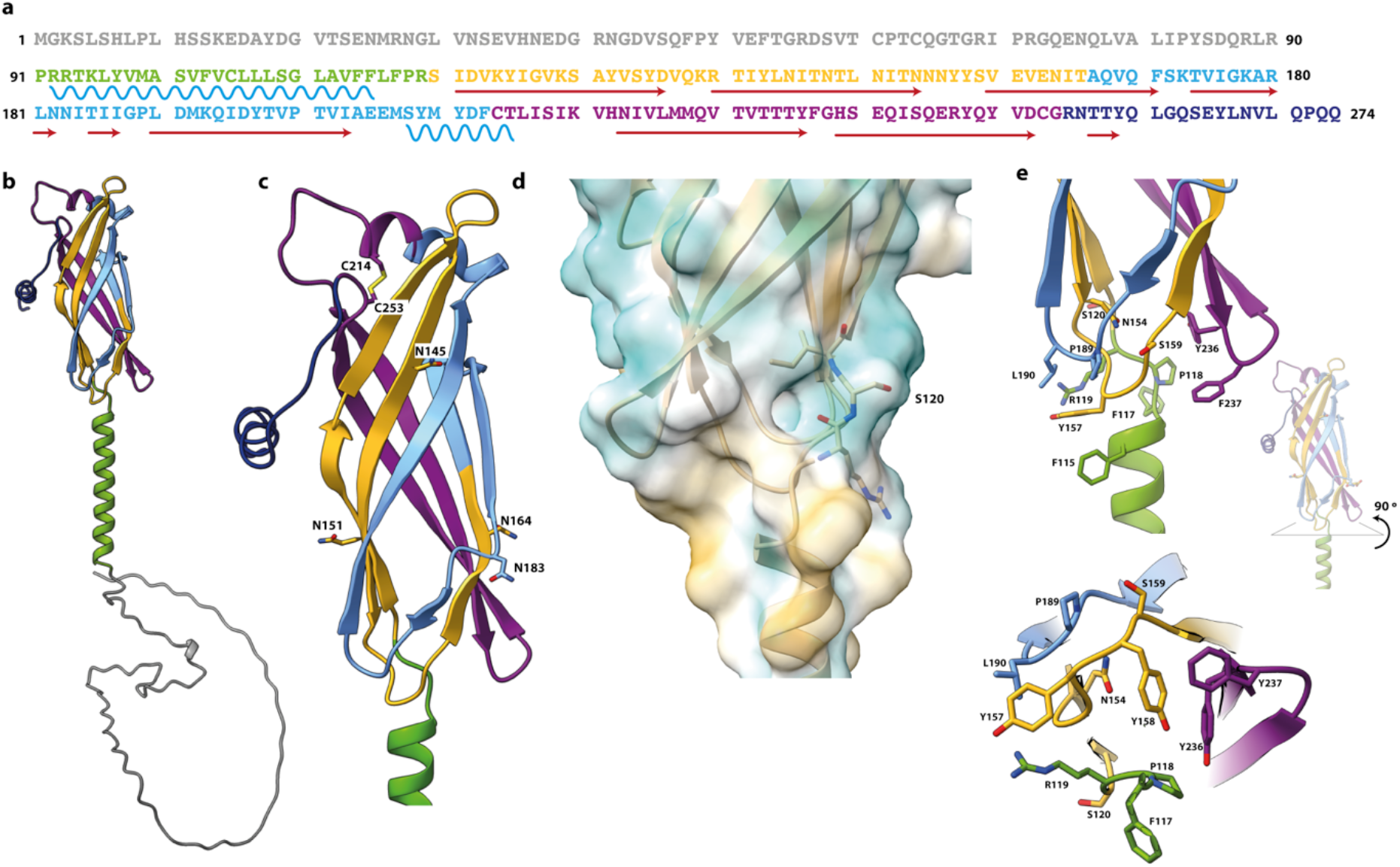
AlphaFold prediction of TMEM106B. **a.** Amino acid sequence of TMEM106B. Predicted α-helices are represented with wavy blue lines; predicted β-strands with red arrows. Residues 1-90, 91-119, 120-166, 167-210, 211-254 and 255-274 are coloured in grey, green, yellow, light blue, magenta and dark blue, respectively. **b.** AlphaFold prediction of TMEM106B. **b.** Close-up view of the AlphaFold prediction of part of the transmembrane helix (green) and the lumenal domain, with glycosylation sites N145, N151, N164, N183 and disulphide bridge C214, C253 shown as sticks. **d.** Hydrophobicity surface view at the interface between the lumenal domain and the transmembrane helix. Residues 119-121 are shown as sticks. **e.** Two orthogonal close-up views of residues close to the lysosomal membrane surface.

**Extended Data Figure 8:**
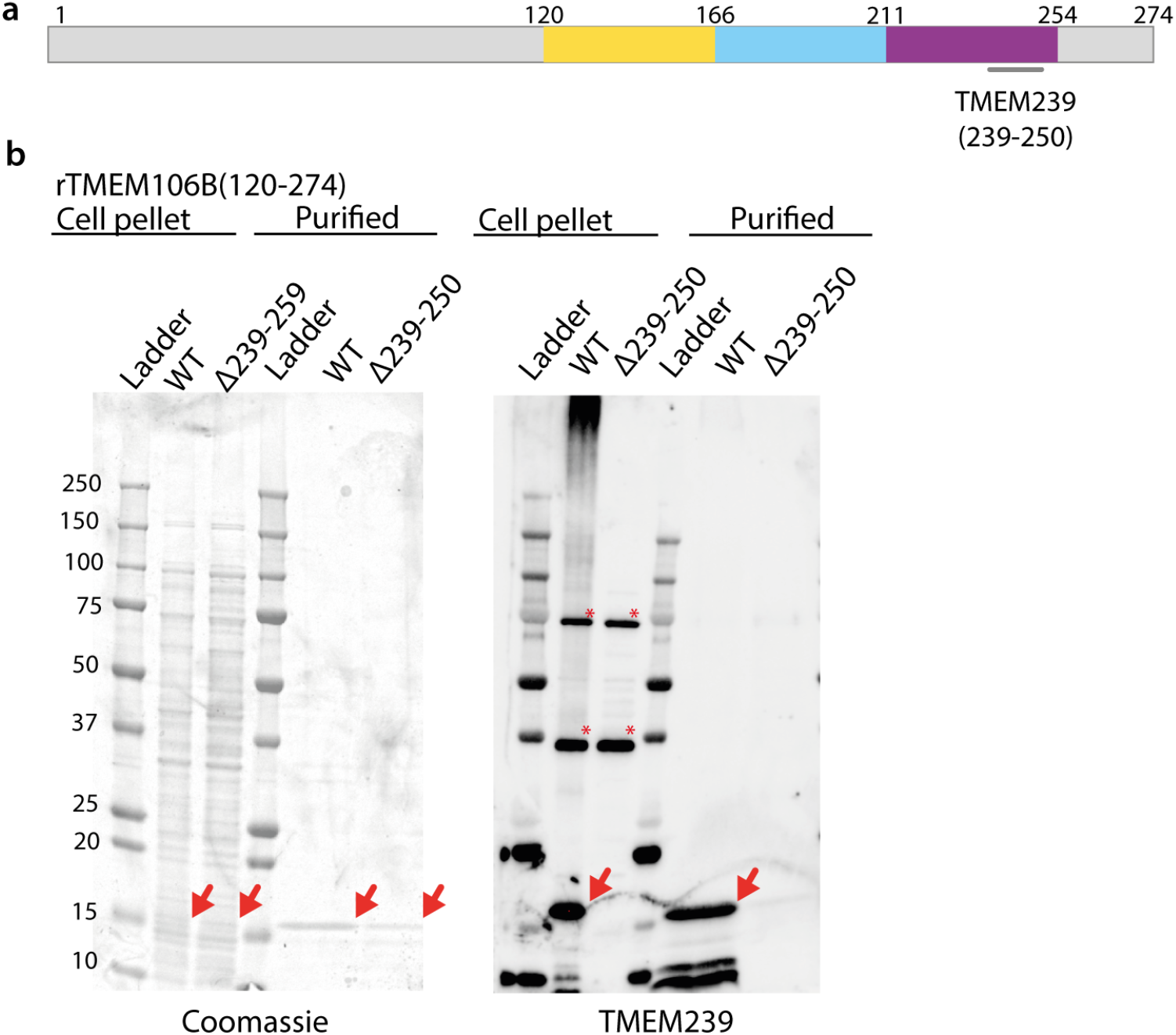
Immunoblot analysis confirms the specificity of antibody TMEM239. **A.** Diagram of the TMEM106B sequence, coloured in accordance with Figure 1, and with the epitope of the TMEM239 antibody (residues 239-250) indicated. **B.** Coomassie stain and immunoblots with recombinant TMEM106B (WT; residues 120-274) and the corresponding epitope-deletion construct (Δ239-250). Red arrows indicate TMEM106B bands; red asterisks indicate non-specific binding.

**Extended Data Table 1:** Further details on cases.

| Case | Disease | Gender | Mutation | Brain region | Publication |
| --- | --- | --- | --- | --- | --- |
| 1 | AD | M | No | FL | Yang et al., 2021: sporadic AD case 1 |
| 2 | FAD | F | *PSEN1* (F105L) | FL | Yang et al., 2021: familial AD case 2 |
| 3 | EOAD | F | No | FL | - |
| 4 | PA | M | No | FL | Yang et al., 2021 |
| 5 | CBD | F | No | FL | Zhang et al., 2020: case 1 |
| 6 | CBD | F | No | FL | Zhang et al., 2020: case 2 |
| 7 | FTDP-17 | M | *MAPT* (P301L) | TL | - |
| 8 | AGD | M | No | NAcc | Shi et al., 2021: AGD case 1 |
| 9 | AGD | M | No | NAcc | Shi et al., 2021: AGD case 2 |
| 10 | LNT | F | No | FL | Shi et al., 2021 |
| 11 | ARTAG | F | No | HPC | Shi et al., 2021; Yang et al., 2021 |
| 12 | PD | M | No | Cg | - |
| 13 | PDD | M | No | AMG | Yang et al., 2021 |
| 14 | FPD | F | *SNCA* (G51D) | TL | - |
| 15 | DLB | M | No | FL | Schweighauser et al., 2020: DLB case 2 |
| 16 | DLB | M | No | FL | Yang et al., 2021 |
| 17 | MSA | F | No | PU | Schweighauser et al., 2020: MSA case 1 |
| 18 | MSA | M | No | PU | Schweighauser et al., 2020: MSA case 5 |
| 19 | MSA | F | No | PU | Schweighauser et al., 2020: MSA case 2 |
| 20 | FTLD-TDP-A | F | *GRN* (intronic) | FL | Yang et al., 2021 |
| 21 | FTLD-TDP-C | F | No | FL | - |
| 22 | ALS | F | No | MC | - |
| 23 | Control | M | No | FL | - |
| 24 | Control | M | No | FL | - |

AD: sporadic Alzheimer's disease; FAD: familial Alzheimer's disease; EOAD: sporadic early-onset Alzheimer's disease; PA: pathological aging; CBD: corticobasal degeneration; FTDP-17T: familial frontotemporal dementia and parkinsonism linked to chromosome 17 caused by *MAPT* mutations; AGD: argyrophilic grain disease; LNT: limbic-predominant neuronal inclusion body 4R tauopathy; ARTAG: aging-related tau astroglipathy; PD: sporadic Parkinson's disease; PDD: sporadic Parkinson's disease dementia; FPD: familial Parkinson's disease; DLB: dementia with Lewy bodies; MSA: multiple system atrophy; FTLD-TDP-A: familial frontotemporal lobar degeneration with TDP-43 inclusions type A; FTLD-TDP-C: sporadic frontotemporal lobar degeneration with TDP-43 inclusions type C; ALS: amyotrophic lateral sclerosis; Control: neurologically normal individual. FL: frontal lobe; TL: temporal lobe; NAcc: nucleus accumbens; HPC: hippocampus; AMG: amygdala; PU: putamen; MC: motor cortex.

**Extended Data Table 2:** Cryo-EM data acquisition and structure determination.

|  | Case 1<br>(AD)<br>Fold I-s | Case 18<br>(MSA)<br>Fold I-d | Case 19<br>(MSA)<br>Fold IIa-s | Case 19<br>(MSA)<br>Fold IIb-s | Case 17<br>(MSA)<br>Fold III-s |
| --- | --- | --- | --- | --- | --- |
| **Data acquisition** |  |  |  |  |  |
| Electron gun | CFEG | XFEG | XFEG | XFEG | XFEG |
| Detector | Falcon 4i | K2 | K2 | K2 | K2 |
| Energy filter slit (eV) | 10 | 20 | 20 | 20 | 20 |
| Magnification | 165,000 | 105,000 | 105,000 | 105,000 | 105,000 |
| Voltage (kV) | 300 | 300 | 300 | 300 | 300 |
| Electron dose (e-/Å<sup>2</sup>) | 40 | 38.8 | 47.5 | 47.5 | 49.2 |
| Defocus range (μm) | 0.6 to 1.4 | 1.8 to 2.4 | 1.7 to 2.6 | 1.7 to 2.8 | 1.7 to 2.8 |
| Pixel size (Å) | 0.727 | 1.15 | 1.15 | 1.15 | 0.829 |
| **Map refinement** |  |  |  |  |  |
| Symmetry imposed | C1 | C2 | C1 | C1 | C1 |
| Initial particle images<br>(no.) | 47,414 | 17,027 | 67,797 | 67,797 | 121,045 |
| Final particle images<br>(no.) | 47,414 | 12,057 | 7,347 | 12,855 | 24,142 |
| Map resolution (Å) | 2.64 | 3.64 | 3.38 | 3.47 | 2.76 |
| FSC threshold | 0.143 | 0.143 | 0.143 | 0.143 | 0.143 |
| Helical twist (°) | -0.42 | -0.41 | -0.63 | -0.49 | -0.69 |
| Helical rise (Å) | 4.81 | 4.78 | 4.78 | 4.79 | 4.79 |
| **Model Refinement** |  |  |  |  |  |
| Model resolution (Å) | 2.6 | 3.74 | 3.22 | 3.33 | 2.41 |
| FSC threshold | 0.5 | 0.5 | 0.5 | 0.5 | 0.5 |
| Map sharpening *B* factor<br>(Å<sup>2</sup>) | -37.57 | -45.37 | -29.29 | -34.66 | -43.93 |
| Model composition |  |  |  |  |  |
| Non-hydrogen atoms | 3426 | 6852 | 3426 | 3426 | 3426 |
| Protein residues | 405 | 810 | 405 | 405 | 405 |
| Ligands | 12 | 24 | 12 | 12 | 12 |
| *B* factors (Å<sup>2</sup>) |  |  |  |  |  |
| Protein | 43.85 | 43.85 | 43.85 | 43.85 | 44.72 |
| Ligand | 51.79 | 51.79 | 51.79 | 51.79 | 52.55 |
| R.m.s. deviations |  |  |  |  |  |
| Bond lengths (Å) | 0.013 | 0.013 | 0.013 | 0.013 | 0.013 |
| Bond angles (°) | 2.086 | 2.086 | 2.149 | 2.286 | 1.943 |
| Validation |  |  |  |  |  |
| MolProbity score | 0.95 | 0.96 | 1.39 | 1.35 | 0.98 |
| Clashscore | 0.00 | 0.15 | 0.88 | 0.59 | 0.00 |
| Poor rotamers (%) | 0.00 | 0.00 | 0.00 | 0.00 | 0.53 |
| Ramachandran plot |  |  |  |  |  |
| Favored (%) | 94.24 | 93.98 | 87.22 | 85.21 | 92.23 |
| Allowed (%) | 5.76 | 6.02 | 12.78 | 14.79 | 7.77 |
| Disallowed (%) | 0.00 | 0.00 | 0.00 | 0.00 | 0.00 |

